# The Treatment of diarrheal mice with Tenebrio Molitor meal

**DOI:** 10.1101/2023.12.21.572900

**Authors:** Tingting Liu, Qiaoli Wang, Zhengli Wang, Jiaxu Yan, Jianjun Zhu, Hong Shen, Jungang Wang

**Affiliations:** College of Animal Science and Technology, Shihezi University,Xinjiang Uygur Autonomous Region, China

**Author notes:** **Corresponding author:** Hong SHEN, Gangjun WANG, **Hong SHEN:** Cellular, **Gangjun WANG:** Cellular. Tingting Liu, Qiaoli Wang, contributed equally to this work. Zhengli Wang, Jiaxu Yan, Jianjun Zhu, Hong Shen, Jungang Wang also contributed equally to this work. **Contact author:** Tingting LIU: Cellular.

**Keywords:** Tenebrio molitor, Mouse, Escherichia coli, Diarrhea, Antioxidant enzymes

## Abstract

Feeding Tenebrio Molitor meal has an important effect on promoting the growth, absorption, reproduction, and disease resistance of animals. In this study, 3×10^8^cfu/ml Escherichia coli was used to establish a mouse diarrhea model. Different doses (8%, 5%, 2.5%) of tenebrio molitor meal were added to the basic diet, respectively. The feed intake, water intake, body weight, loose stool rate, diarrhea rate, intestinal flora number, immune organ index, intestinal enzyme, and serum enzyme activities of the diarrhea mice were detected. The results showed that compared with the model group, the feed intake, water intake, and body weight of mice with diarrhea were improved by adding tenebrio molitor meal, and the dosage was proportional to that of tenebrio molitor meal. The rate of loose stool and diarrhea decreased with the increase of tenebrio molitor meal. The total bacterial count and Escherichia coli count in the intestinal tract of mice with diarrhea were negatively correlated with the dosage of tenebrio molitor meal. The immune organ index of the diarrhea mice in the three tenebrio molitor meal supplementation groups was higher than that in the model group and was proportional to the dosage. The liver index of the 8% tenebrio molitor meal supplementation group was 11.79mg/g higher than that in the blank group. Compared with the blank group, diarrhea significantly decreased the activities of various enzymes in the intestinal tract and serum of mice (P<0.05). Compared with the model group, the activities of intestinal and serum protective enzymes (superoxide dismutase, peroxidase, catalase), detoxification enzymes (glutathione-S transferase, acetylcholinesterase, acid phosphatase) and digestive enzymes (serum amylase, serum lipase, lactate dehydrogenase) in diarrhea mice were increased by adding tenebrio molitor meal (P<0.05). The results showed that tenebrio molitor meal had a positive effect on the treatment of diarrhea in mice.

## Introduction

Diarrhea is a major disease that harms humans and young animals. To prevent and control the infection of the agents of the disease, a large number of antibiotics are used, but due to the irrational use of antibiotics, the agents have resistance ^[1]^. The diarrhea rate and mortality of children and young animals caused by E. coli infection are extremely high, which can reach 10%-30%^[2]^. How to reduce the diarrhea rate caused by E. coli has become an urgent problem to be solved. Tenebrio molitor, also known as tenebrio molitors, belong to the genus Coleoptera. First found in South America, it is a kind of world grain storage pest with the characteristics of fast reproduction and easy feeding^[3]^. Tenebrio molitor is rich in active substances protein, amino acid, fatty acid, and fiber, which can be used as a renewable animal protein resource to alleviate the shortage of animal feed resources^[4]^. Buβler et al. found that tenebrio molitor can be eaten directly^[5]^. As a medicinal resource, tenebrio molitor flavus has made some achievements. Studies have found that adding tenebrio molitor meal to the basic diet of obese mice can change the expression of glucose and lipid metabolism genes in their liver and adipose tissue^[6]^. Khanal et al.^[7]^ found that fee ding tenebrio molitor meal could increase animal weight and promote the digestion and absorption of protein. Biasato et al.^[8]^ found that feed ingtenebrio molitor can improve the influence of serum acidic and alkaline enzyme activity of animals, thus enhancing the antioxidant capacity of animals. At present, the experiments on the treatment of E. coli diarrhea by tenebrio molitor mainly focus on the medicinal activity of tenebrio molitor in vivo, and the studies on tenebrio molitor meal are very few. As a sustainable drug resource, the medicinal value of tenebrio molitor has been reported in the fields of anti-inflammation, diarrhea, tumors, and other diseases^[9]^, but whether its insect powder has such effects has not been reported.

In this study, three doses (8%, 5%, 2.5%) of tenebrio molitor meal were added to the basic diet of mice with diarrhea to study the therapeutic mechanism of tenebrio molitor meal on mice with diarrhea and provide a basis for clinical application. To strengthen the cultivation scale and technical research of medicinal resources is to better serve our human society with this sustainable and valuable resource.

### Test method

In the early stage, according to the experimental method of LuJunjia et al.^[10]^ 3×10^8^cfu/ml Escherichia coli (ATCC43255) was selected and the mouse diarrhea model was constructed by injecting bacterial suspension into the abdomen.

### Animals

Death by pentobarbital anesthesia and cervical dislocation (Grab the tail of the mice with the right hand and pull it back hard, while the left thumb and index finger press down on the head of the mice, pull off the spinal cord and brain, and the mice will die immediately. Mice were monitored daily for signs of pain or distress according to the guidelines of the Animal Experimentation Ethics Committee of the School of Animal Science and Technology, Shihezi University.

### Ethics statement

The study was conducted strictly in accordance with the university in an already approved facility. All mice were reared and euthanized in strict accordance with the guidelines of the Animal Experiment Ethics Committee of the College of Animal Science and Technology, Shihezi University, approval number: 2021-0021.

### Prepare tenebrio molitor meal feed

Experimental basic feed (wheat 60%, corn 25%, pea 25%, soybean meal 10%, cabbage 15%, apple 15%, hawthorn 15%, sweet potato 2 0%, celery 15%, lettuce 10%) was purchased from the experimental animal breeding center of Shihezi University. The tenebrio molitor was killed at a low temperature (-4°C), and the surface debris was washed off with running water. It was dried to constant weight in an incubator at 55°C, crushed, and mixed with the basic feed, and the dosage of tenebrio molitor meal was 8%, 5%, and 2.5%, respectively.

### Mice were grouped and treated

A total of 75 healthy male Kunming mice with a body weight of (20±2)g were randomly divided into 5 groups with 3 replicates per group and 5 mice per replicate. On the 1st, 2nd, 3rd, 4th, and 5th days, 3×10^8^cfu/ml Escherichia coli (ATCC43255) bacterial suspension was injected intraperitoneally, the dose was 0.02ml/g, and the diarrhea model was established. The blank group was injected with an equal dose of sterile normal saline. After the diarrhea model was built, the diarrhea group was divided into model groups, 8% tenebrio molitor meal group, 5% tenebrio molitor meal group, and 2.5% tenebrio molitor meal group. During treatment, the model group and blank group were fed a basic diet without tenebrio molitor meal. The experiment lasted for 20 days, including 5 days of adaptation period, 5 days of diarrhea model period, and 10 days of feeding tenebrio molitor meal treatment period. The mice were managed according to routine feeding^[11]^, with a feed dosage of 55g/ cage and water of 150 ml/ cage to ensure adequate supply.

### Body weight, feed intake, and water intake were measured

The feeding situation of the mice was observed every day, and the cage was regularly fed and cleaned. The weight, feed intake, and water intake of the mice were respectively weighed at the 1d, 3d, 5d, 7d, 9d, 11d, 13d, and 15d (16:00-18:00), and the average weight, feed intake, and water intake of the mice were calculated.

Individual feed intake = (amount of fed food - amount of remaining food) /5

Single drinking water quantity = (water supply - remaining water) /5

### The rates of loose stool and diarrhea were measured

The shape of mouse feces was determined by reference to previous methods^[12]^. Observe and record the number of diarrhea and the number of loose stools in mice at 1, 3, 5, 7, 9, 11, 13, and 15d, and calculate the rate of loose stools and diarrhea in animals, the formula is as follows:

Loose stools rate = Number of loose stools/total animal defecation ×100%

Diarrhea rate = Number of animals with diarrhea/total number of animals ×100%

### The rates of loose stool and diarrhea were measured

The shape of mouse feces was determined by reference to previous methods ^[12]^. Observe and record the number of diarrhea and the number of loose stools in mice at 1, 3, 5, 7, 9, 11, 13, and 15d, and calculate the rate of loose stools and diarrhea in animals, the formula is as follows:

Loose stools rate = Number of loose stools/total animal defecation ×100%

Diarrhea rate = Number of animals with diarrhea/total number of animals ×100%

### Determination of the number of intestinal colonies in mice

On the 20th day of the experiment, all mice were killed by spinal dislocation. The contents of the ileum, cecum, and colon of each group were collected in a sterile EP tube and placed in a refrigerator at -80°C for the determination of total bacteria and E. coli in the ileum, cecum, and colon. Escherichia coli and total bacteria were cultured using eosin methylene blue medium and AGAR medium respectively, and the determination steps and methods were referred to previous test methods^[13]^.

### Determination of immune organ index in mice

After treatment (20d), the mice were weighed, their liver, spleen, and thymus were taken, the surfaces were washed with 4°C normal saline, and the surface water was absorbed by filter paper and weighed. The indexes of the liver, spleen, and thymus were calculated with the following formula:

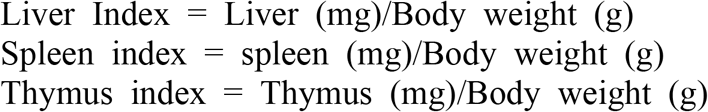

### Determination of intestinal enzyme activity in mice

At the end of the experiment (20d), the ileum, cecum, and colon tissues of mice were collected, their surfaces and intestinal contents were washed with sterile normal saline at 4°C, placed into 1.5ml sterile EP tube, and stored at -20°C for later use. Weigh and grind the ileum, cecum, and colon tissues of 0.2g each in a mortar, pour in liquid nitrogen and grind, add 1.8mlsterile normal saline at 4°C to make tissue homogenate, centrifuge for 10min at 4000r/min, and take the supernatant into a 2ml sterile EP tube at 4°C. Superoxide dismutase (SOD), per oxidase (POD), catalase (CAT), glutathione-S-transferase (GSH), acid phosphatase (ACP), acetylcholinesterase (ACHe), amylase (AMS) and fat in ileum, cecum and colon were determined by ELISA kit (Shanghai Yucun Biotechnology Co., LTD.) Enzyme (LPS), lactate dehydrogenase (LDH) activity.

### Determination of serum enzyme activity in mice

At the end of the experiment (20d), a small amount of ether was used to stun mice, and blood was collected through the heart with a 2 mL syringe. The blood was placed in a 2mL sterile EP tube, centrifuged at 4000r/min for 10min immediately, and serum was collected and placed in a refrigerator at 4°C. It was used to determine the activities of superoxide dismutase (SOD), peroxidase (POD), catalase (CAT), glutathione-S-transferase (GSH), acid phosphatase (ACP), acetylcholinesterase (ACHe), amylase (AMS), lipase (LPS) and lactate dehydrogenase (LDH) in serum.

### Data statistics and analysis

The test data were analyzed by SPSS17.0, One-WayANOVA.

## Results and analysis

### Effects of tenebrio molitor meal on body weight, feed intake, and water intake of diarrhea mice

During the treatment of diarrhea mice with tenebrio molitor meal, (Table 1) on day 7, the body weight of mice in 8% and 2.5% tenebrio molitor meal group was 21.17g and 20.65g, respectively, significantly lower than that in the blank group (24.60g) (P < 0.05). At 9, 11, and 13 days, the body weight of the 2.5% tenebrio molitor meal group was significantly lower than that of the blank group (P < 0.05), but there was no significant difference between each dose group and the model group. On day 15, the body weight of mice in the 8%, 5% tenebrio molitor meal group, and blank group was 27.59g, 28.42g, and 29.54g, respectively, which were higher than 24.24g in the model group. In conclusion, dietary tenebrio molitor meal has a positive effect on the weight increase of diarrhea mice. (Table 2) On day 11, the feed intake of mice in 8%, 5%, and 2.5% tenebrio molitor meal groups was 29.38g, 31.29g, and 30.91g, respectively, which was significantly lower than that in the blank group at 41.40g (P < 0.05) and higher than that in model group at 22.41g. The feed intake of the 8% tenebrio molitor meal group was lower than that of the 5% and 2.5% tenebrio molitor meal group. On day 13, the feed intake of the 2.5% group was lower than that of the blank group (P < 0.05). On day 15, the feed intake of 8%, 5%, and 2.5% groups was 34.92g, 29.26g, and 25.65g, respectively, which was significantly lower than that of the blank group by 41.06g (P < 0.05), and significantly higher than that of the model group by 21.24g. The difference between the blank group and the model group was 19.82g, indicating that the feed intake of mice with diarrhea caused by E. coli was decreased. (Table 3) Compared with the model group, an increase in the water consumption of diarrhea mice was observed in the three groups supplemented with tenebrio molitor meal, and the water consumption of 8% tenebrio molitor meal group was higher than that of other treatment groups; On day 13, the drinking water volume of 2.5% tenebrio molitor meal group and model group was 10.25mL and 10.50mL, respectively, which was significantly lower than that of blank group (P < 0.05); On day 15, the water intake of 8% and 5% tenebrio molitor meal groups and blank group was 23.00mL, 23.00mL and 28.50mL, respectively, which were significantly higher than 12.75mL in model group (P < 0.05).

**Table 1.**
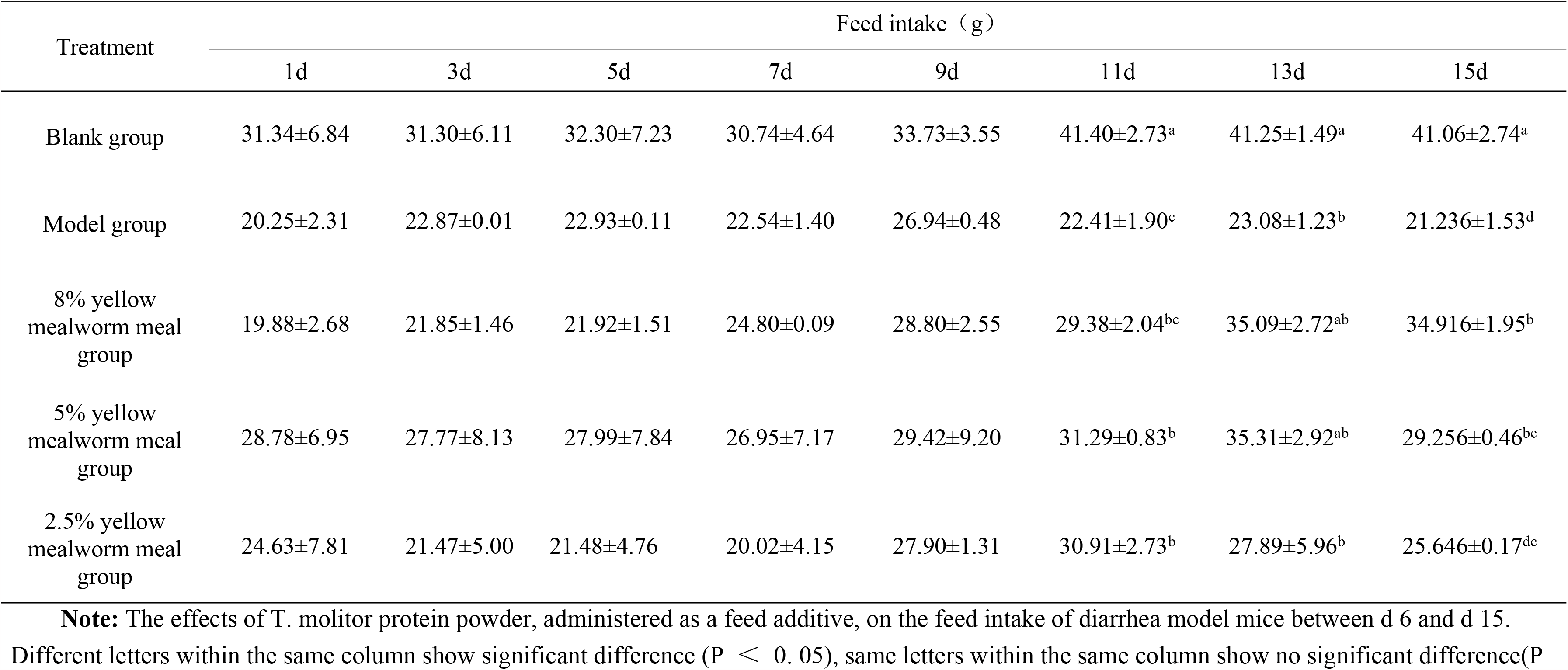
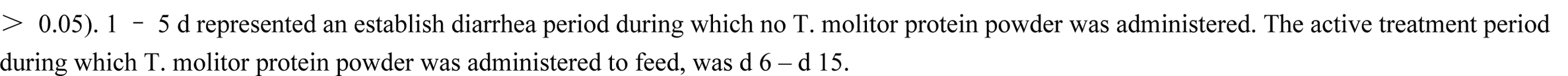
Effected of Tenebrio molitor powder on feed intake of diarrhea mice.

| Treatment | Feed intake (g) |  |  |  |  |  |  |  |
| --- | --- | --- | --- | --- | --- | --- | --- | --- |
|  | 1d | 3d | 5d | 7d | 9d | 11d | 13d | 15d |
| Blank group | 31.34±6.84 | 31.30±6.11 | 32.30±7.23 | 30.74±4.64 | 33.73±3.55 | 41.40±2.73<sup>a</sup> | 41.25±1.49<sup>a</sup> | 41.06±2.74<sup>a</sup> |
| Model group | 20.25±2.31 | 22.87±0.01 | 22.93±0.11 | 22.54±1.40 | 26.94±0.48 | 22.41±1.90<sup>c</sup> | 23.08±1.23<sup>b</sup> | 21.23±1.53<sup>d</sup> |
| 8% yellow mealworm meal group | 19.88±2.68 | 21.85±1.46 | 21.92±1.51 | 24.80±0.09 | 28.80±2.55 | 29.38±2.04<sup>bc</sup> | 35.09±2.72<sup>ab</sup> | 34.91±1.95<sup>b</sup> |
| 5% yellow mealworm meal group | 28.78±6.95 | 27.77±8.13 | 27.99±7.84 | 26.95±7.17 | 29.42±9.20 | 31.29±0.83<sup>b</sup> | 35.31±2.92<sup>ab</sup> | 29.25±0.46<sup>bc</sup> |
| 2.5% yellow mealworm meal group | 24.63±7.81 | 21.47±5.00 | 21.48±4.76 | 20.02±4.15 | 27.90±1.31 | 30.91±2.73<sup>b</sup> | 27.89±5.96<sup>b</sup> | 25.64±0.17<sup>dc</sup> |

**Note:** The effects of *T. molitor* protein powder, administered as a feed additive, on the feed intake of diarrhea model mice between d 6 and d 15. Different letters within the same column show significant difference ( $P < 0.05$ ), same letters within the same column show no significant difference ( $P > 0.05$ ).
> 0.05). 1 – 5 d represented an establish diarrhea period during which no T. molitor protein powder was administered. The active treatment period during which T. molitor protein powder was administered to feed, was d 6 – d 15.

**Table 2.** Effected of Tenebrio molitor powder on water intake of diarrhea mice.

| Treatment | Water intake (mL) |  |  |  |  |  |  |  |
| --- | --- | --- | --- | --- | --- | --- | --- | --- |
|  | 1d | 3d | 5d | 7d | 9d | 11d | 13d | 15d |
| Blank group | 11.50±5.5 | 13.50±0.50 | 14.50±0.50 | 11.50±0.50 | 14.50±0.50 | 30.00±1.00 | 28.00±2.00<sup>a</sup> | 28.50±3.50<sup>a</sup> |
| Model group | 3.00±0.00 | 9.00±0.00 | 10.00±0.00 | 15.00±2.00 | 18.00±4.00 | 12.75±0.25 | 10.50±0.50<sup>b</sup> | 12.75±0.25<sup>b</sup> |
| 8% yellow mealworm meal group | 3.50±1.50 | 12.75±0.25 | 11.50±0.50 | 14.00±3.00 | 19.50±6.50 | 20.00±6.00 | 19.50±5.50<sup>ab</sup> | 23.00±3.00<sup>a</sup> |
| 5% yellow mealworm meal group | 3.50±1.50 | 10.00±1.00 | 10.00±1.00 | 11.00±2.00 | 15.75±3.25 | 18.00±6.00 | 18.00±5.00<sup>ab</sup> | 23.00±1.00<sup>a</sup> |
| 2.5% yellow mealworm meal group | 3.50±0.50 | 7.50±3.50 | 9.75±2.75 | 11.75±1.25 | 14.25±0.75 | 12.50±5.00 | 10.25±2.25<sup>b</sup> | 14.50±0.50<sup>b</sup> |

**Note:** The effects of T. molitor protein powder, administered as a feed additive, on the body weight of diarrhea model mice between d 6 and d 15. Different letters within the same column show significant difference ( $P < 0.05$ ), same letters within the same column show no significant difference ( $P > 0.05$ ). 1 – 5d represented an establish diarrhea period during which no T. molitor protein powder was administered. The active treatment period during which T. molitor protein powder was administered to feed, was 6 – 15d.

**Table 3.** Effected of Tenebrio molitor powder on body weight of diarrhea mice.

| Body weight |
| --- |

| Treatment | 1d | 3d | 5d | 7d | 9d | 11d | 13d | 15d |
| --- | --- | --- | --- | --- | --- | --- | --- | --- |
| Blank group | 19.81±0.50 | 22.03±0.56 | 23.22±0.47<sup>a</sup> | 24.60±0.40<sup>a</sup> | 26.08±0.51<sup>a</sup> | 27.24±0.27<sup>a</sup> | 28.58±0.27<sup>a</sup> | 29.54±0.35<sup>a</sup> |
| Model group | 20.29±0.59 | 20.61±0.46 | 21.02±0.33<sup>ab</sup> | 22.12±0.36<sup>ab</sup> | 23.54±0.42<sup>ab</sup> | 24.59±0.60<sup>ab</sup> | 24.53±0.52<sup>b</sup> | 24.23±0.74<sup>c</sup> |
| 8% yellow mealworm meal group | 18.97±0.21 | 19.42±0.36 | 19.66±0.38<sup>ab</sup> | 21.17±0.66<sup>b</sup> | 23.62±1.02<sup>ab</sup> | 25.35±0.72<sup>ab</sup> | 26.77±0.70<sup>ab</sup> | 27.59±0.48<sup>ab</sup> |
| 5% yellow mealworm meal group | 20.85±0.74 | 21.66±1.10 | 21.82±0.89<sup>ab</sup> | 22.42±0.95<sup>ab</sup> | 25.47±1.10<sup>a</sup> | 26.51±1.19<sup>a</sup> | 28.00±1.12<sup>a</sup> | 28.42±1.46<sup>a</sup> |
| 2.5% yellow mealworm meal group | 20.23±0.61 | 20.87±0.63 | 21.52±0.75<sup>ab</sup> | 20.65±0.41<sup>b</sup> | 21.76±0.84<sup>b</sup> | 22.67±0.75<sup>b</sup> | 24.53±0.44<sup>b</sup> | 25.41±0.34<sup>bc</sup> |

**Note:** The effects of *T. molitor* protein powder, administered as a feed additive, on the body weight of diarrhea model mice between d 6 and d 15. Different letters within the same column show significant difference ( $P < 0.05$ ), same letters within the same column show no significant difference ( $P > 0.05$ ). 1 – 5d represented an establish diarrhea period during which no *T. molitor* protein powder was administered. The active treatment period during which *T. molitor* protein powder was administered to feed, was 6 – 15d.

**Table 4.** Effected of Tenebrio molitor powder on diarrhea rate of diarrhea mice.

| Treatment | Diarrhea rate (%) |  |  |  |  |  |  |  |
| --- | --- | --- | --- | --- | --- | --- | --- | --- |
|  | 1d | 3d | 5d | 7d | 9d | 11d | 13d | 15d |
| Blank group | 0 | 0 | 0 | 0 | 0 | 0 | 0 | 0 |
| Model group | 20.00 | 60.00 | 92.00 | 90.00 | 80.00 | 80.00 | 80.00 | 80.00 |
| 8% yellow mealworm meal group | 20.00 | 73.00 | 85.00 | 60.00 | 50.00 | 30.00 | 20.00 | 20.00 |
| 5% yellow mealworm meal group | 33.00 | 60.00 | 93.00 | 70.00 | 60.00 | 50.00 | 50.00 | 40.00 |
| 2.5% yellow mealworm meal group | 20.00 | 20.00 | 87.00 | 80.00 | 70.00 | 60.00 | 60.00 | 50.00 |

**Note:** The effects of *T. molitor* protein powder, administered as a feed additive, on the diarrhea rate of diarrhea model mice between d 6 and d 15. 1 – 5d represented an establish diarrhea period during which no *T. molitor* protein powder was administered. The active treatment period during which *T. molitor* protein powder was administered to feed, was d 6 – d 15.

**Table 5.** Effect of Tenebrio molitor powder on <u>stool</u> rate of diarrhea mice.

| Treatment | <u>Stool</u> rate |  |  |  |  |  |  |  |
| --- | --- | --- | --- | --- | --- | --- | --- | --- |
|  | 1d | 3d | 5d | 7d | 9d | 11d | 13d | 15d |
| Blank group | 0 | 0 | 0 | 0 | 0 | 0 | 0 | 0 |
| Model group | 69.23 | 51.72 | 54.84 | 62.5 | 48.15 | 51.72 | 56.67 | 64.29 |
| 8% yellow mealworm meal group | 46.67 | 50.00 | 57.14 | 40.63 | 40.74 | 36.36 | 29.16 | 16.67 |
| 5% yellow mealworm meal group | 40.00 | 51.85 | 59.26 | 48.39 | 44.83 | 35.00 | 30.00 | 23.08 |
| 2.5% yellow mealworm meal group | 40.00 | 56.00 | 57.69 | 51.72 | 50.00 | 45.45 | 38.09 | 38.09 |

**Note:** The effects of *T. molitor* protein powder, administered as a feed additive, on the stool rate of diarrhea model mice between d 6 to 15d. 1 – 5d represented an establish diarrhea period during which no *T. molitor* protein powder was administered. The active treatment period during which *T. molitor* protein powder was administered to feed, was d 6 –15d.

These results indicate that adding tenebrio molitor meal to the diet of diarrhea mice can significantly increase their body weight, feed intake, and water intake.

### Effects of tenebrio molitor meal on the number of intestinal Escherichia coli and total bacteria in diarrhea mice

The balance of intestinal flora has a positive significance for the stable operation of the body. (Figure 1) The number of Escherichia coli in the model group was significantly higher than that in the blank group and the three tenebrio molitor meal addition groups. The maximum number of Escherichia coli in the cecal model group was 220.9×10^6^CFU/mL, which was 2.9 times that of the blank group. The number of Escherichia coli in the ileum, colon, and cecum of 8% tenebrio molitor meal supplementation group had no significant difference compared with the blank group, but the number of Escherichia coli among the three supplementation groups decreased with the increase of tenebrio molitor meal dosage. In conclusion, the tenebrio molitor meal has a certain inhibitory effect on the production of enteric Escherichia coli in mice with diarrhea. (Figure 2) The total bacterial count of the model group was higher than that of the other groups, and the total bacterial count of the cecum model group was the largest, up to 620.6×106CFU/mL, while the total bacterial count of the 8% tenebrio molitor meal group of the ileum was at least 28.6×104CFU/mL. Except for 2.5% tenebrio molitor meal in the cecum group, the total bacteria number of each dose group in the ileum, colon, and cecum was significantly lower than that in the corresponding model group, and the total bacteria number of each dose group decreased with the increase of tenebrio molitor meal supplemental level. In conclusion, dietary tenebrio molitor meal can inhibit the increase of intestinal total bacteria number in diarrhea mice to a certain extent.

**Fig1.**
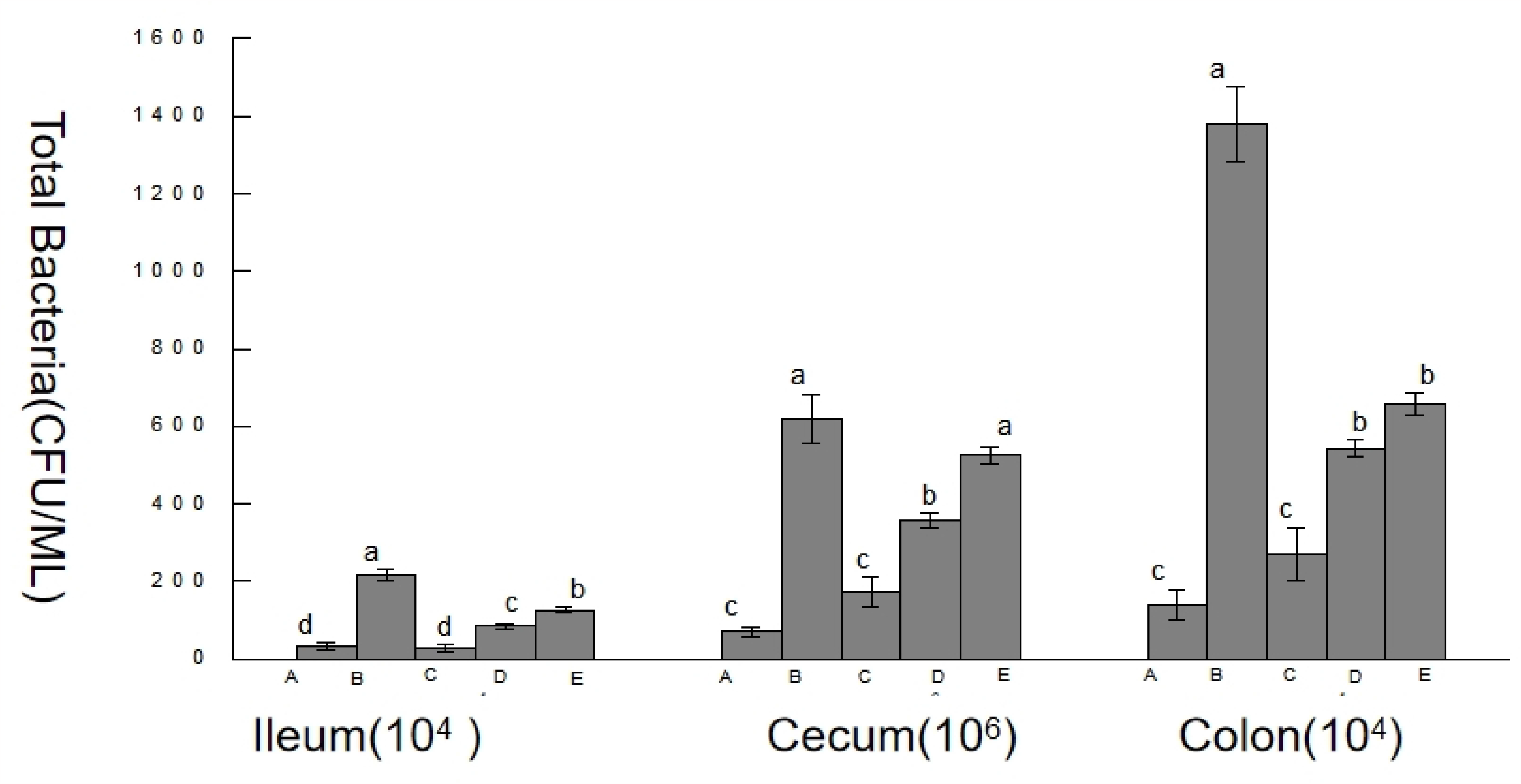
Effected of Tenebrio molitor powder on Escherichia coli of ileum, colon, cecum to diarrhea mice. **Note:** The diluted with aseptic saline contents of the ileum, cecum, and colon were diluted with 10^4^, 10^5^, 10^6^ CFU/mL gradients. It was used to count the number of colonies of Escherichia coli in intestinal contents. The was the optimum count number of colonies in 30-300. The was 10^4^ cfu/mL concentration of ileum and colon content, and the final concentration of cecum was 10^6^ cfu/mL, the lower case letters showed significant difference. A, B, C, D and E respectively indicate that the blank group, the model group, the 8% dose group, the 5% dose group, the 2.5% dose group, different small letters showed significant difference (*p* <0.05), same small letters showed no significant difference(*p>*0.05).

**Fig2.**
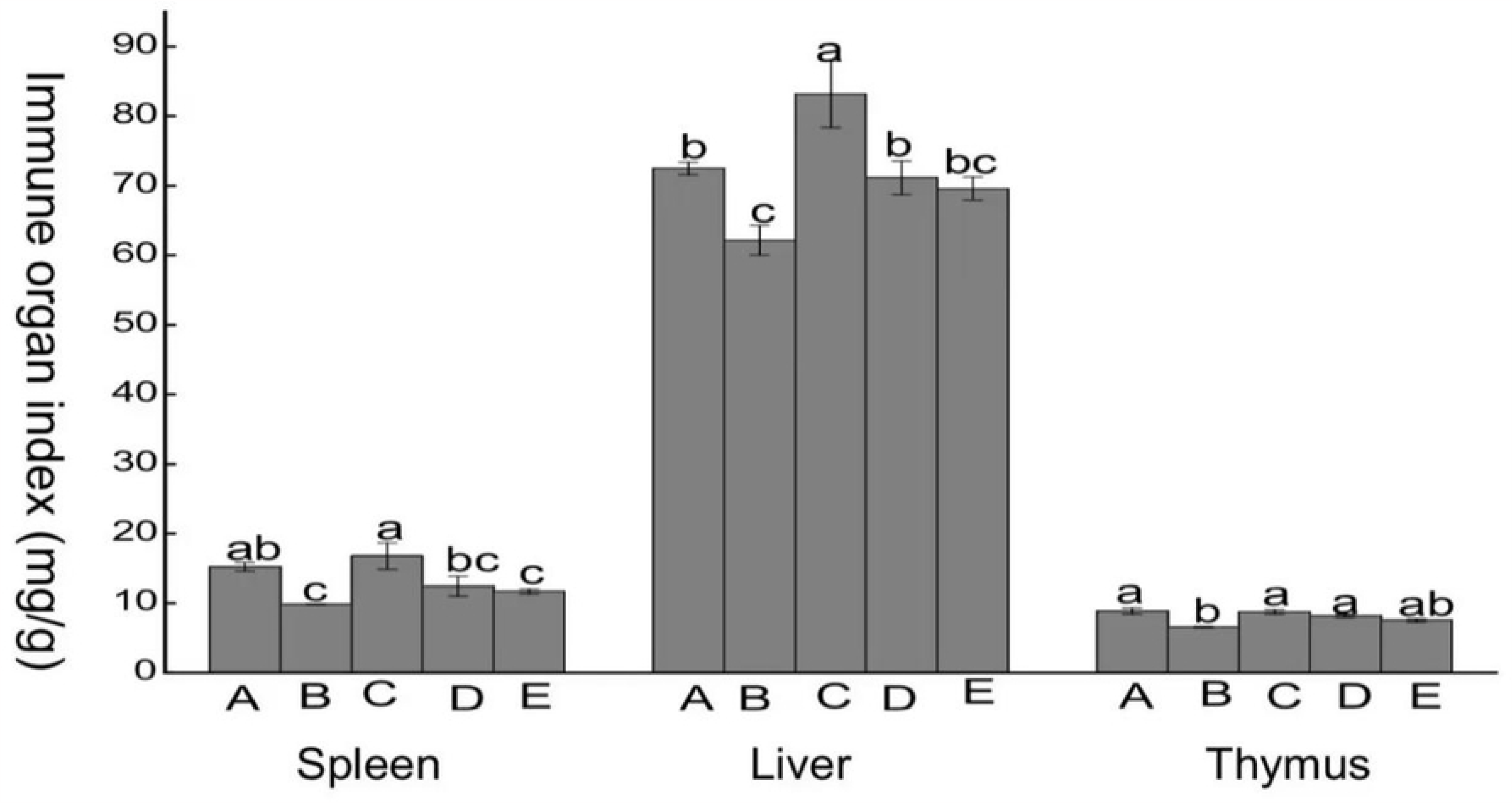
Effected of Tenebrio molitor powder on the number of total bacteria of ileum, colon, cecum to diarrhea mice. **Note:** The diluted with aseptic saline contents of the ileum, cecum, and colon were diluted with 104, 105, 106 cfu/mL gradients. It was used to count the number of colonies of total bacterial in intestinal contents. The was the optimum count number of colonies in 30-300. The was 104 cfu/mL concentration of ileum and colon content, and the final concentration of cecum was 106 cfu/mL, the lower case letters showed significant difference. A, B, C, D and E respectively indicate that the blank group, the model group, the 8% dose group, the 5% dose group, the 2.5% dose group, different small letters showed significant difference (p <0.05), same small letters showed no significant difference (p>0.05).

### Effect of tenebrio molitor meal on immune organs of diarrhea mice

(FIG. 3) The highest spleen index of the 8% tenebrio molitor meal group was 11.79mg/g, which was significantly different from the model group (P < 0.05) and increased by 1.57mg/g compared with the blank group. The liver index of the 8% tenebrio molitor meal group was the highest, reaching 78.16mg/g, which was significantly higher than that of other groups (P < 0.05). The thymus index of the model group was the smallest, only 1.57mg /g, which was significantly lower than that of 8%, 5% tenebrio molitor meal groups, and blank group (P < 0.05). In conclusion, dietary tenebrio molitor meal can improve the immune organ index of mice with diarrhea, and there is a positive correlation with the supplemental amount.

**Fig3.**
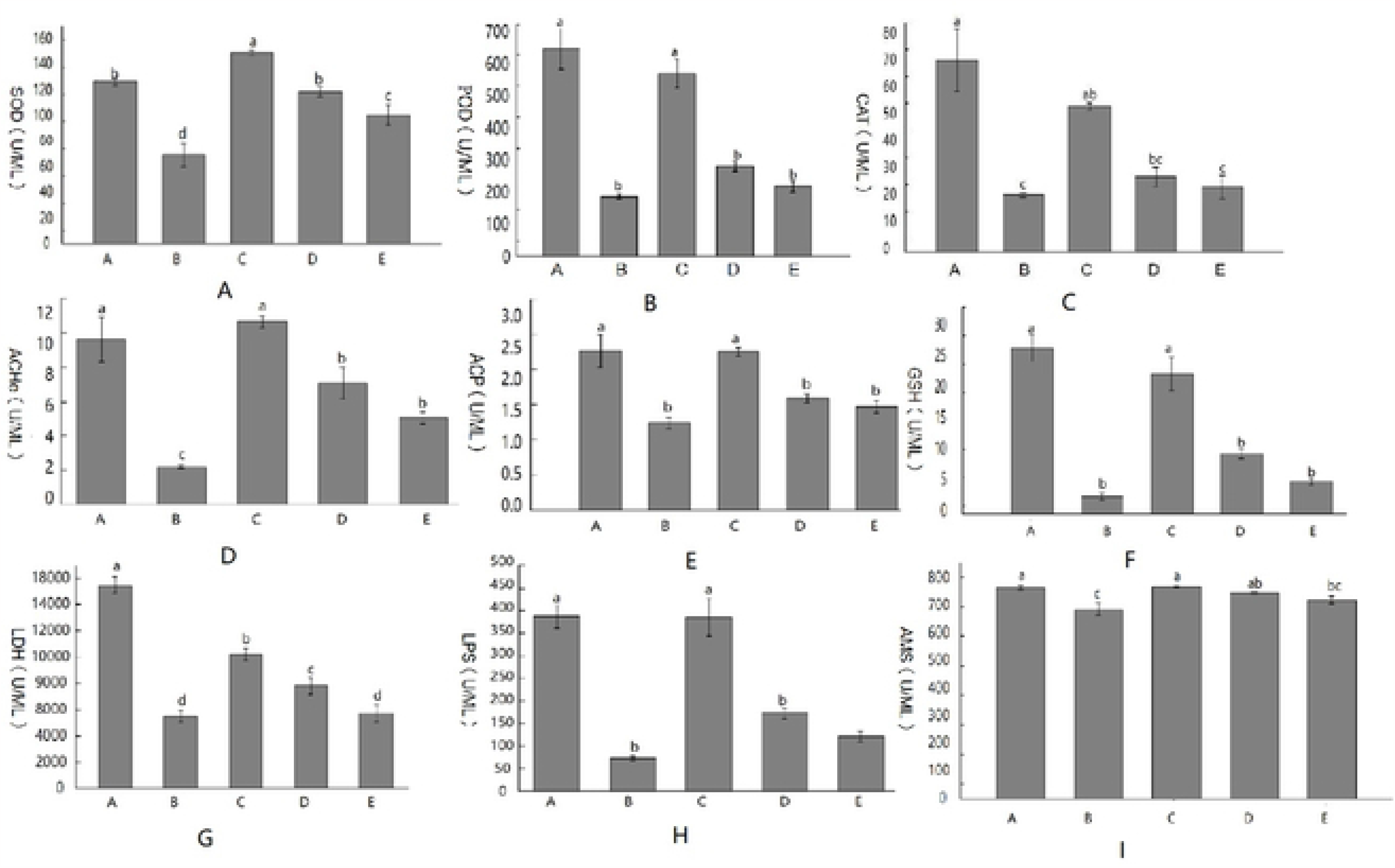
Effect of Tenebrio molitor powder on immune organs of diarrhea mice. **Note:** the lower case letters showed significant difference. different small letters showed significant difference (p <0.05), same small letters showed no significant difference (p>0.05), A, B, C, D and E respectively indicate that the blank group, the model group, the 8% dose group, the 5% dose group, the 2.5% dose group.

### Effect of tenebrio molitor meal on intestinal enzyme activity in mice with diarrhea

Enzyme activity in the intestinal tract of mice with diarrhea was measured by an ELISA kit (Shanghai Yuchun Biotechnology Co, LTD.). The results showed that the contents of protective enzymes (superoxide dismutase, peroxidase, catalase), detoxification enzymes (glutathione-S transferase, acetylcholinesterase, acid phosphatase), and digestive enzymes (serum amylase, serum lipase, lactate dehydrogenase) in the ileum of mice caused by E. coli decreased. It produces a large number of free radicals in the colon, cecum, and ileum. (FIG. 4A) The highest superoxide dismutase (SOD) activity was 609 .67U/mL in the 8% tenebrio molitor group. The lowest value was 86.51U/mL in the ileum model group. The activity of SOD in the ileum, colon, and cecum of diarrhea mice was increased in the three tenebrio molitor meal groups. The effects of different doses of tenebrio molitor meal on enzyme activity were also different. It can be seen that the 8% tenebrio molitor meal supplementation group had the most obvious improvement in the SOD activity of diarrhea mice, and the activity was close to that of the blank group. (FIG. 4B) The highest POD activity was 96.25U/mL in the blank cecum group, while the lowest POD activity was only 1.84U/mL in the ileal model group. Compared with the model group, the POD enzyme activity of the ileum, colon, and cecum of diarrhea mice was increased by adding tenebrio molitor meal, and was proportional to adding tenebrio molitor meal. The POD enzyme activity of the 8% tenebrio molitor meal added to the group was the highest among the three added groups and close to the blank control group. (Figure 4C) The highest CAT activity was 59.23U/mL in the blank group of cecum, and the lowest CAT activity was 4.4U/mL in the ileal model group. Compared with the model group, the three tenebrio molitor meal supplementation groups could increase the intestinal CAT enzyme activity of diarrhea mice. The CAT activity of 8% tenebrio molitor meal in the colon and cecum was very close to that of the blank control group. These results indicate that the CAT activities of the ileum, cecum, and colon of diarrhea mice can be significantly increased by dietary tenebrio molitor meal. (Figure 4D) The highest ACHe content in the cecum was 4.41U/mL, and the lowest ACHe content in the ileum was 0.24U/mL. Compared with the model group, the increase of ACHe enzyme activity was observed in the three groups supplemented with tenebrio molitor meal. The increase in enzyme activity was proportional to the amount of tenebrio molitor meal added. The ACHe activity of the 8% tenebrio molitor meal supplementation group was higher than that of the blank control group. The data show that adding the proper amount of tenebrio molitor meal to the diet of mice with diarrhea has a positive effect on the activity of intestinal ACHe. (Figure 4E) The highest ACP content in the cecum was 816.77U/mL, while the lowest ACP content in the colon was 134.07U/mL. Compared with the model group, the ACP content of the three dose groups was significantly increased and was proportional to the addition amount. Among them, the ACP activity of the ileum and cecum in the 8% tenebrio molitor meal supplementation group was close to that in the blank control group. (Figure 4F) The highest GSH activity was 1076.93U/mL in the blank cecum group. GSH activity in the ileum was the lowest, only 199.67U/mL. Compared with the model group, the GSH content of the three groups supplemented with tenebrio molitor meal was significantly increased and was proportional to the additional amount. The GSH activity of 8% tenebrio molitor meal was closest to that of the blank control group, and the enzyme activity of 5% and 2.5% tenebrio molitor meal in the ileum was close to that of the control group. (FIG. 4G) LDH activity in the blank cecum group was the highest, reaching 7737.05U/mL. LDH activity in the ileum model group was the lowest, only 708.1U/mL. The LDH content of the three added tenebrio molitor meal groups was higher than that of the model group and was proportional to the added amount. The content of LDH in the ileum was not affected by the tenebrio molitor meal between different dosage groups. (FIG. 4H) The LPS activity in the blank cecum group was the highest, reaching 871.6U/mL. LPS activity in the ileum model group was the lowest, only 50.86U/mL. Compared with the model group, the content of intestinal LPS of diarrhea mice was significantly increased in the three dose groups. The content of LPS in the ileum of the 8% tenebrio molitor meal supplementation group was slightly higher than that in the control group. The results indicated that tenebrio molitor meal played a positive role in enhancing intestinal enzyme activity in mice with diarrhea. (FIG. 4I) The AMS activity in the blank caecum group was the highest, reaching 1. 13U/mL. AMS activity in the ileum model group was the lowest (0.33U/mL). The activity of AMS in the added tenebrio molitor meal group was higher than that in the model group and was proportional to the added amount. The content of AMS in the ileum of the 8% tenebrio molitor meal supple mentation group was slightly higher than that of the control group. tenebrio molitor meal can improve intestinal enzyme activity and intestinal antioxidant capacity in mice with diarrhea.

**Fig4.**
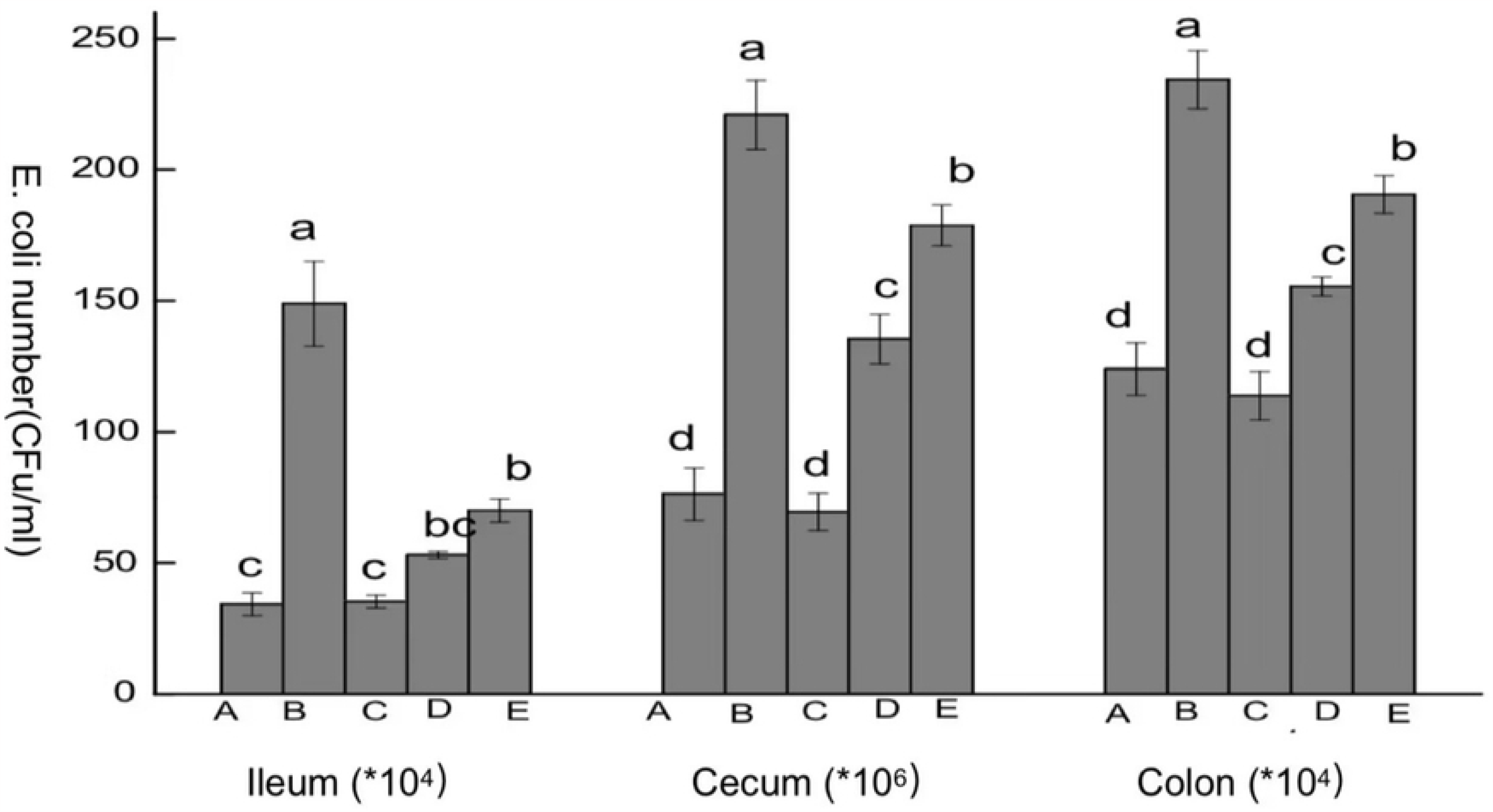
Effect of tenebrio molitor meal on various enzyme activities in ileum, cecum and colon of mice. **Note:** A, B, C, D and E respectively indicate that the blank group, the model group, the 8% dose group, the 5% dose group, the 2.5% dose group, different small letters showed significant difference (p <0.05), same small letters showed no significant difference (p>0.05).

### Effect of tenebrio molitor meal on serum enzyme activity in mice with diarrhea

The level of serum enzyme activity reflected the antioxidant activity of serum. ELISA kit was used to determine the activity of various enzymes in the serum of mice with diarrhea. (FIG. 5A) The highest superoxide dismutase (SOD) activity in the 8% tenebrio molitor meal group was 140.7U/mL, which was 80.13U/mL higher than that in the model group. The activity of serum superoxide dismutase (SOD) in the blank group and three dose groups was significantly higher than that in the model group (P < 0.05). The serum SOD activity of the 8% tenebrio molitor group was significantly higher than that of the blank group (P < 0.05). The serum SOD activity of the blank group and 5% tenebrio molitor meal group was significantly higher than that of 2.5% tenebrio molitor meal group (P < 0.05). tenebrio molitor meal can increase the activity of superoxide dismutase (SOD) in the serum of mice with diarrhea. (Figure 5B) The peroxidase (POD) activity in the blank group was the highest (670.37U/mL), and the peroxidase (POD) activity in the model group was the lowest (193.52U/mL). Compared with the model group, the three tenebrio molitor meal groups can increase the activity of serum catalase, and it is proportional to the additional amount. Serum POD activity in the blank group and 8% tenebrio molitor meal group was significantly higher than that in 5% and 2.5% tenebrio molitor meal groups (P < 0.05). (Figure 5C) With the increase of tenebrio molitor meal supplemental level, the serum catalase (CAT) activity of diarrhea mice also increased. The highest serum catalase (CAT) activity in the blank group was 70.94U/mL, which was 49.82U/mL higher than 21.12U/mL in the model group. Serum catalase (CAT) activity in the 8% tenebrio molitor meal group and blank group was significantly higher than that in the model group and 2.5% tenebrio molitor meal group (P < 0.05). (FIG. 5D) Serum glutathione-S transferase (GS H) activity in the blank group was 30.34U/mL, 26.27U/mL higher than that in a model group of 4.07U/mL. The serum glutathione-S transferase (GSH) activity of the blank group and 8% tenebrio molitor meal group was significantly higher than that of 5% and 2.5% tenebrio molitor meal groups and model groups (P < 0.05). The activity of serum glutathione-S transferase (GSH) in mice with diarrhea could be increased by adding tenebrio molitor meal. (Figure 5E) Serum acid phosphatase (ACP) activity in the blank group was 2.26U/mL, 1.02U/mL higher than 1.24U/mL in the model group. The activity of serum acid phosphatase (ACP) in mice with diarrhea could be increased by different doses of tenebrio molitor meal. The activity of serum acid phosphatase (ACP) in the blank group and 8% tenebrio molitor meal group was significantly higher than that in 5% and 2.5% tenebrio molitor meal groups and model groups (P < 0.05). (FIG. 5F) The highest serum acetylcholinesterase (ACHe) activity in the 8% tenebrio molitor meal group was 10. 63/mL, which was 8.43U/mL higher than that in model group of 2.2U/mL. The serum acetylcholinesterase (ACHe) activity in the blank group and 8% tenebrio molitor meal group was significantly higher than that in the 5%, 2.5% tenebrio molitor meal group and model group (P < 0.05). (FIG. 5G) Serum lactate dehydrogenase (LDH) activity in the blank group was 15477.66/mL, which was 9972.51U/mL higher than that in model group 5505.15U/mL . Different doses of tenebrio molitor could increase serum lactate dehydrogenase activity in mice with diarrhea. The serum lactate dehydrogenase (LDH) activity of 8% and 5% tenebrio molitor meal groups and blank group was significantly higher than that of the model group (P < 0.05). (FIG. 5H) Serum lipase (LPS) activity in the blank group was 387.08U/mL, which was 314.03U/mL higher than 73.05U/mL in the model group. The activity of serum lipase (LPS) in the serum of mice with diarrhea could be increased by different doses of tenebrio molitor meal. The serum lipase activity of the blank group and 8% tenebrio molitor meal group was significantly higher than that of the 5%, 2.5% tenebrio molitor meal group and model group (P < 0.05). (FIG. 5I) Serum amylase (AMS) activity of the 8% tenebrio molitor meal group was 717.49U/mL, which was 76.51U/mL higher than that of the model group 640.98U/mL. The activity of serum amylase (AMS) in the serum of mice with diarrhea could be increased by different doses of tenebrio molitor. Serum lipase activity in the 8% tenebrio molitor meal group and blank group was significantly higher than that in the model group (P < 0.05). Adding tenebrio molitor meal to the basal diet can increase the activity of the serum enzyme system and increase the serum antioxidant capacity of diarrhea mice.

**Fig 5.**
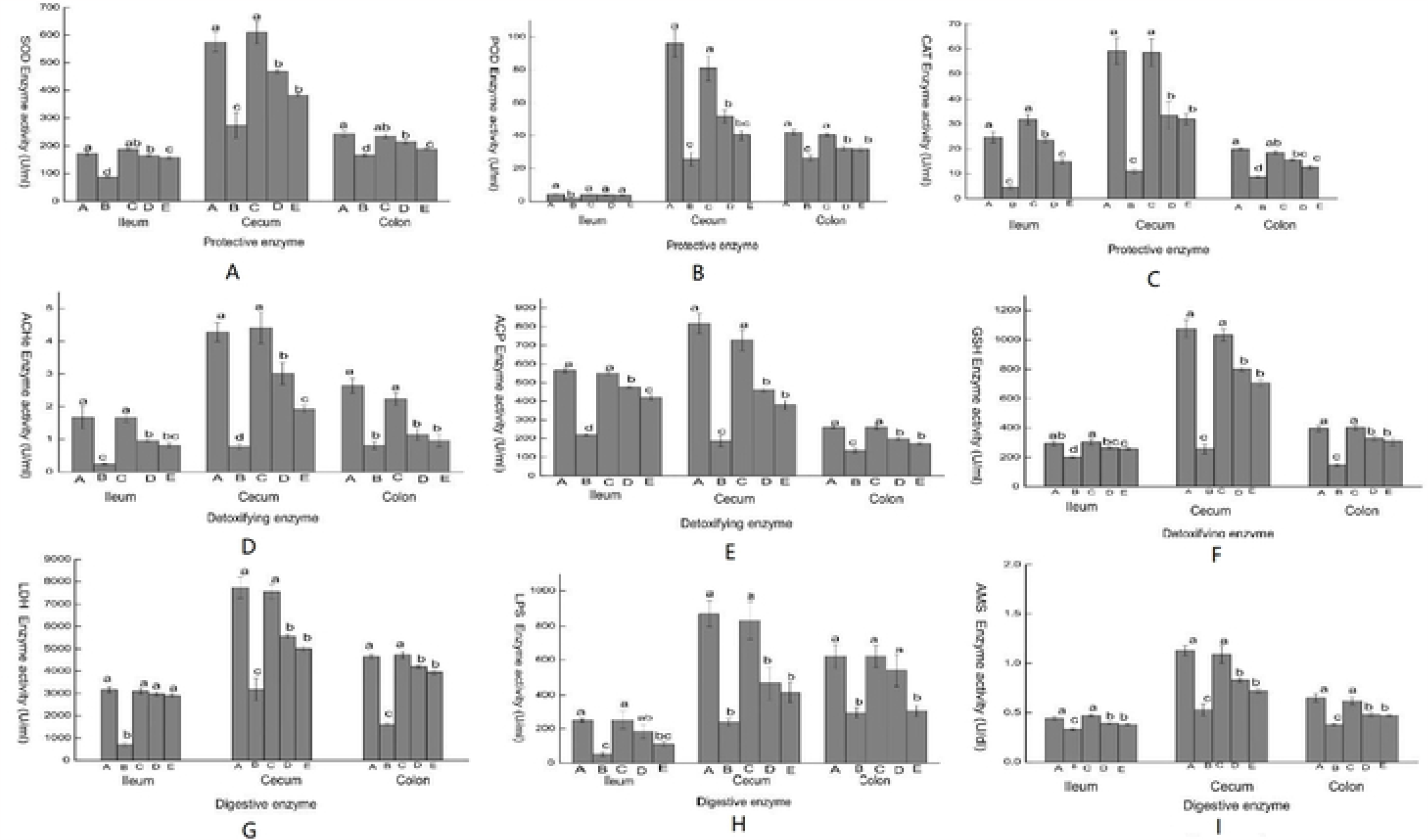
Effects of mealworm meal on the activities of various enzymes in mouse serum. **Note:** A, B, C, D and E respectively indicate that the blank group, the model group, the 8% dose group, the 5% dose group, the 2.5% dose group, different small letters showed significant difference (p <0.05), same small letters showed no significant difference (p>0.05).

## Discuss

Escherichia coli is one of the major pathogens causing diarrhea in humans and animals ^[14]^. The diarrhea rate and mortality of animals caused by E. coli infection are extremely high, which can reach 10%-30%^[15]^. Escherichia coli contains adhesion and enterotoxin, which is the main factor causing diarrhea^[16]^. Enterotoxin promotes intestinal tissue lesions, causes intestinal environment disorders, increases the volume of loose stools, and thus causes diarrhea and death in mice ^[17]^. Tao Wu et al. made a diarrhea model by gavage and intraperitoneal injection of E. coli ^[18]^ and found that an ideal diarrhea model with a high diarrhea rate and low mortality could be obtained after intraperitoneal injection of E. coli. Based on this, Escherichia coli (ATC C43255) was selected in the experiment, and the mouse diarrhea model was constructed by intraperitoneal injection of bacterial suspension.

Relevant studies have shown that the reasons for the differences in body weight, feed intake, and water consumption of mice may be related to the different needs of different individuals for food and water consumption, the preparation process of tenebrio molitor meal, and the bioactive ingredients contained in tenebrio molitor meal ^[19-20]^. The basic diet of the 8% tenebrio molitor meal group is rich in amino acids, proteins, unsaturated fatty acids, and other nutrients ^[21]^, which is conducive to the absorption of water and food by mice, resulting in weight gain of mice. The nutrient content of 5% and 2.5% tenebrio molitor meal groups was relatively low. This is similar to the results of Momin et al., which found that fly maggot meal can promo tedigestion, immunity, and growth of animals, but there are some differences, which may be related to the difference between feeding worm meal and experimental subjects^[22]^.

The mouse gut usually contains a stable and complex microbial community that plays important roles in the body, including enhancing the accumulation of host fat and short-chain fatty acids and regulating the mucosal immune system^[23]^. Studies have shown that most additives have the function of regulating the stability of the intestinal microbial community by affecting the number of intestinal bacteria^[24]^. Various antibacterial active substances contained in tenebrio molitors are related, such as antibacterial substance MD-(7095), chitose, chitosan, antibacterial protein, etc^.[25]^. This is consistent with our findings that each dose group can significantly reduce the diarrhea rate, loose stool rate, total bacteria, and Escherichia coli number in the intestinal tract (ileum, colon, cecum) of mice with diarrhea.

The difference in the liver index of mice may be related to the different dosages of tenebrio molitor meal. With the increase in dosage, fat was deposited in the liver, increasing liver index ^[26]^. The reason for the difference in spleen and thymus indices of mice among different dosage groups of tenebrio molitor meal may be that the tenebrio molitor meal added to the feed can improve the immunity of mice with diarrhea, and the spleen and thymus, as part of the immune system, also improve with the increase of tenebrio molitor meal added dose.

Superoxide dismutase (SOD) has the function of anti-oxidation and anti-aging, and its mechanism is mainly to clear the harmful superoxide anion free radical (O2-). The level of SOD content indirectly reflects the ability of the body to remove superoxide anion (O2-) ^[27]^. Peroxidase (POD) is the hallmark enzyme of peroxisome, which mainly exists in the peroxisome of carrier, and can remove toxic substances with high activity in the body and reduce cell damage^[28]^. Catalase (CAT) can accelerate the decomposition of peroxides in the body, to avoid the damage of oxides in the machine^[29]^. Glutathione-s transferase (GSH) plays an important role in preventing the oxidative decomposition of hemoglobin, maintaining the activity of sulfhydryl protein, and ensuring the reducibility of sulfhydryl protein and the integrity of cells ^[30]^. Acid phosphatase (ACP) can catalyze the hydrolysis of almost all monophosphate esters and directly participate in phosphorus metabolism, playing an important role in the digestion, absorption, and secretion of calcium and phosphorus^[31]^. Acetylcholinesterase (ACHe) can degrade acetylcholine, ensure the normal transmission of nerve signals in organisms, participate in c ell development and maturation, and promote neuronal development and nerve regeneration ^[32]^. Lactate dehydrogenase (LDH) is involved in intracellular REDOX reactions that convert the resulting lactic acid to pyruvate. The level of LDH is often used to assess the degree of cell damage or inflammatory response ^[33]^. Serum amylase (AMS), which plays an important role in the digestion of polysaccharide compounds in food, is mainly used in the diagnosis of acute pancreatitis ^[34]^. The synergistic effect of these enzymes enables the body to operate in a balanced and stable manner. In this study, diarrhea caused a significant decrease in serum and intestinal enzyme system activity in mice compared with a blank control group. Compared with the model group, tenebrio molitor meal could significantly increase the activity of the enzyme system in the serum and intestinal tract of mice with diarrhea. The effect of different dosages of tenebrio molitor meal on the activity of the enzyme system is also different, which is proportional to the added amount. The results showed that the tenebrio molitor meal could improve the intestinal enzyme activity by increasing the activities of detoxification enzymes (GSH, ACHe, ACP), protective enzymes (SOD, POD, CAT), digestive enzymes (AMS, LPS, LDH), and enhance the anti-diarrhea ability of mice. However, the underlying biological mechanism needs further study, and it is speculated that there is a certain relationship with the chemical substances contained in the body of tenebrio molitors.

The results of this study showed that adding tenebrio molitor meal to the basic diet of mice played a positive role in the treatment of diarrhea in mice. It laid a foundation for developing the medicinal methods and efficacy of tenebrio molitor, better utilizing its effective and healthy medical information, and further promoting the research and development of tenebrio molitor medicinal resources in the prevention and control of human and animal infectious diseases.

## Data Availability

The data reported here have been deposited at the Dryad with the website https://datadryad.org/stash/share/vHLN1lr8z1DpXkt6_LMi0gKtgBBjATCDWjF2NKtvgvM. The DOI number is DOI:10.5061/dryad.xgxd254p7.

## Notes

### Competing Interest Statement

The authors have declared no competing interest.

